# Activation-induced re-organization of chromatin in human T cells

**DOI:** 10.1101/2020.06.08.135020

**Authors:** Naiara G. Bediaga, Hannah D. Coughlan, Timothy M. Johanson, Alexandra L. Garnham, Gaetano Naselli, Jan Schröder, Liam G. Fearnley, Esther Bandala-Sanchez, Rhys S. Allan, Gordon K. Smyth, Leonard C. Harrison.

**Affiliations:** The Walter and Eliza Hall Institute of Medical Research, Parkville, 3052, Australia; Department of Medical Biology, The University of Melbourne, Parkville, 3010, Australia; School of Mathematics and Statistics, The University of Melbourne, Parkville, 3010, Australia

**Keywords:** Human, T cell, activation, chromatin, accessibility, remodelling, genomic interactions, TAD, gene expression

## Abstract

Remodelling of chromatin architecture is known to regulate gene expression and has been well characterized in cell lineage development but less so in response to cell perturbation. Activation of T cells, which triggers extensive changes in transcriptional programs, serves as an instructive model to elucidate how changes in genome organization orchestrate gene expression in response to cell perturbation. To characterize coordinate changes at different levels of chromatin architecture, we analysed chromatin accessibility, chromosome conformation and gene expression after activation of human T cells. T cell activation led to widespread changes in chromatin interactions and accessibility that were mostly shared between CD4^+^ and CD8^+^ T cells. Differential chromatin interactions were associated with upregulation or downregulation of linked target genes. Moreover, activation was associated with the formation of shorter chromatin interactions, partitioning of topologically associating domains (TADs) and acquisition of new TAD boundaries characterized by higher nucleosome occupancy, and lower chromatin accessibility and gene expression. These findings render an integrated and multiscale characterization of activation-induced re-organization of chromatin architecture underlying gene transcription in human T cells.

## BACKGROUND

Mammalian genomes are folded into highly organized hierarchical structures linked to function at each level ^[1]^. At the primary level of chromatin structure, the nucleosome, a 147 base-pair DNA segment wrapped around an octamer of histone proteins, directly influences gene expression by dictating access of DNA to the transcriptional machinery ^[2–4]^. At an intermediate level, the genome is organized into protein-mediated loops that facilitate interaction between of pairs of genomic sites such as promoters and enhancers distant along the linear genome. At a higher level, the genome is organized into self-interacting chromatin, called topologically associating domains (TADs) ^[5,6]^. Disruption of TAD boundaries has been associated with developmental defects ^[7]^ but the functional significance of changes in TAD architecture are otherwise largely unknown. Further, chromosomes are organized into a gene-rich, transcriptionally active compartment (A) with open chromatin and active histone marks and a gene-poor, transcriptionally inactive compartment (B) with condensed chromatin and gene silencing histone marks. Within this overall organization, the interplay between chromatin structure ^[8]^ and gene expression ^[9–11]^ is cell-specific and mediated by transcription factors (TFs) and other DNA binding proteins the functions of which depend on chromatin accessibility.

The immune system evolved to respond to environmental stimuli and exhibits a high degree of phenotypic and functional plasticity in response to external cues. T cells have a central role in the adaptive immune system and are activated under different conditions to expand and differentiate into a variety of specialized, functionally distinct subsets. T cell activation is likely to be instructive of how coordinate changes in chromatin structure orchestrate gene expression programs that underlie rapid and often lifesaving responses. It was recently shown that T cell activation is associated with marked reorganization of chromatin structure at the level of accessibility (measured by ATAC-seq)^[12–14]^ and, separately, conformation (measured by promoter capture-Hi-C) ^[15]^. The present study is the first to analyse genome-wide chromosome conformation in conjunction with chromatin accessibility and whole transcriptome expression, to understand how coordinated changes in higher- and lower-order chromatin structures are linked to gene expression in response to T cell activation.

## RESULTS

### Chromatin accessibility and 3D genomic interactions are immune cell-type specific

Naïve and activated CD4+ T cells, naïve and activated CD8+ T cells, and mature B cells were isolated from two healthy human male donors. Each cell lineage from each donor was profiled using assay for transposase-accessible chromatin using sequencing (ATAC-seq) to generate genome-wide maps of chromatin accessibility, in situ Hi-C to interrogate 3D genomic interactions and RNA-seq for whole transcriptome expression. The consistent use of two independent biological replicates across all three genomic technologies was considered an important aspect of the study, and all bioinformatics analyses used statistically rigorous methods that assessed statistical significance between the cell populations relative to biological variability between the individual donors.

ATAC-seq, Hi-C-seq and RNA-seq sequencing read densities at genes that define the cell phenotypes confirmed specificity for each cell type (Fig. 1A, 1B, 1C). We observed a strong enrichment of ATAC-seq reads at transcription start sites (TSSs) genome-wide, including promoter-TSSs of the cell type-specific expressed genes *CD4, CD8A, IFNG, BCL11B and MS4A1*, an indication of the quality of the data ^[16]^ (Fig. 1A-B). T cells, but not B cells, exhibited robust enrichment of Hi-C genomic interactions at the BCL11B gene, known to be expressed in T but not B cells ^[17]^ (Fig. 1C). Conversely, B cells exhibited robust enrichment of Hi-C genomic interactions at the BCL11A gene, known to be expressed in B but not T cells (Fig. S1). Unsupervised clustering of the samples by multidimensional scaling of the chromatin accessibility, chromatin interaction and gene expression data demonstrated distinct chromatin structure and gene expression signatures for different cell types (Fig. 1D), in keeping with cell-type specificity of chromatin architecture ^[8]^. Samples clustered by activation status over dimension 1 and diverged by cell lineage over dimension 2. By calculating the proportion of variation explained by each dimension using the Glimma package (v1.6.0) ^[18]^ we observed that activation (dimension 1) accounted for the largest proportion of variation in chromatin accessibility, chromatin interactions and gene expression (34%, 27% and 49%, respectively), followed by lineage (dimension 2) (23%, 19% and 17%) (Fig. 1D). Consistent with the smaller contribution of T cell lineage to biological variance in chromatin architecture and interactions, direct comparison of activated CD4^+^ and CD8^+^ T cells showed only a small number of chromatin architecture differences, viz. 864 differentially accessible regions and 69 differential interactions, indicating that activation has a similar effect on chromatin remodelling in CD4^+^ and CD8^+^ T cells.

**Figure 1.**
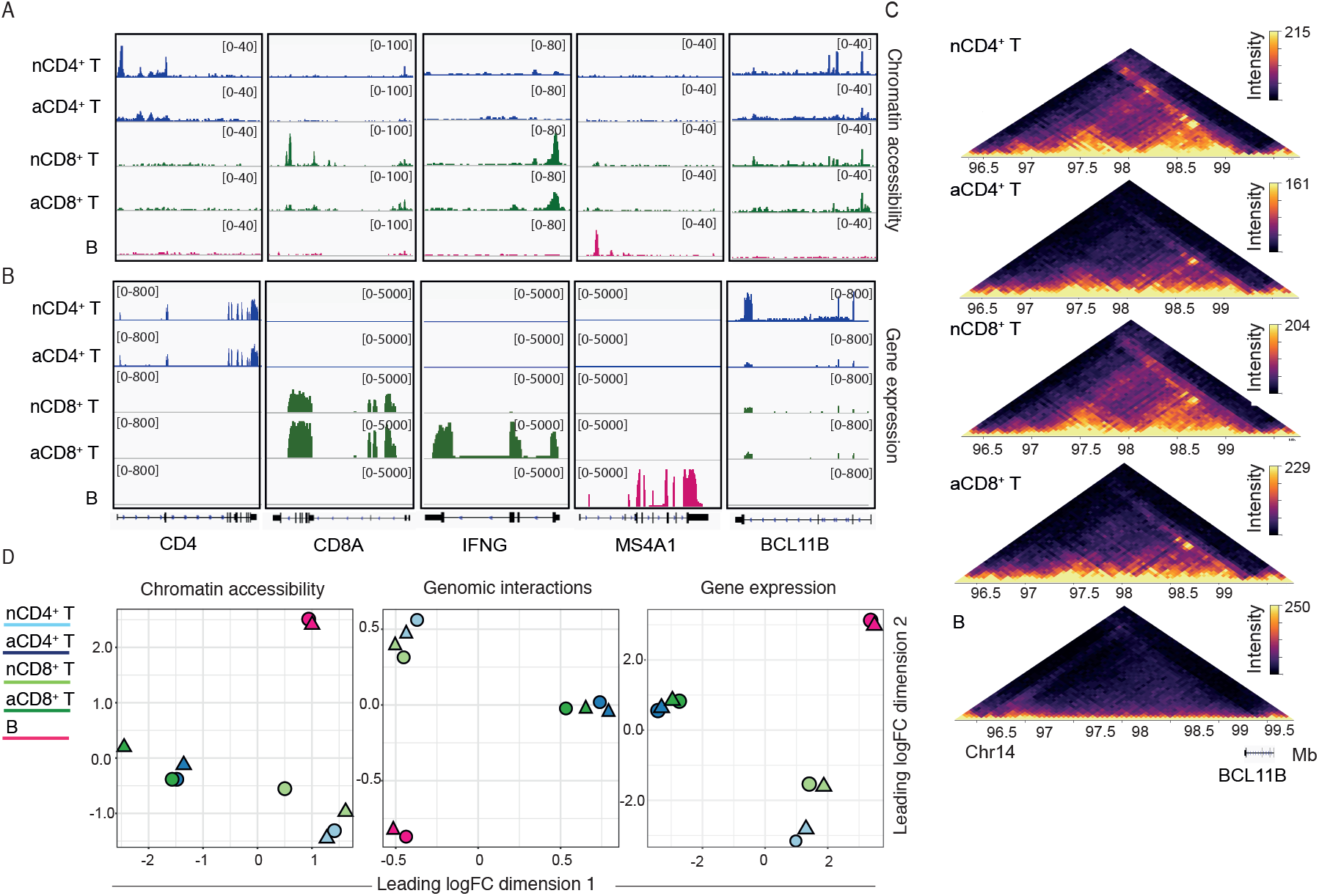
Chromatin structure and gene expression are immune cell type-specific. Normalized read coverage plots of (A) ATAC-seq and (B) RNA-seq libraries at phenotype-defining genes including *CD4*, *CD8A, IFNG, MS4A1* and *BCL11B* loci in resting CD4^+^ (nCD4^+^), resting CD8^+^ (nCD8^+^), activated CD4^+^ (aCD4^+^) and activated CD8^+^ (nCD8^+^) T cells. (C) *In-*situ Hi-C contact matrices for a 1 Mb region on chromosome 14 that includes BCL11B, which is known to be expressed in T but not B cells. The top four Hi-C matrices display data from resting and activated T cells, while the bottom Hi-C matrix shows data from B cells. Color scale indicates number of reads per bin pair. (D) Multidimenional scaling (MDS) plots of log-CPM values over dimensions 1 and 2 with samples coloured by groups and shaped by donors. The log-CPM values were corrected for the donor variable. Distances on the plot correspond to the leading fold-change, which is the average (root-mean-square) log2-fold-change for the 500, 5,000 and 50,000 genes (RNA-seq), peaks (ATAC-seq) or bins (Hi-C) most divergent between each pair of samples.

### T cell activation leads to widespread changes in chromatin accessibility

Activation increased accessibility of 8,484 (7.5 %) and 7,883 (6.9 %) ATAC-peaks in CD4^+^ and CD8^+^ T cells, respectively, while 2,961 (2.6 %) and 3,184 (2.8 %) peaks lost accessibility (Fig. 2A, Table S1). Compared to accessibility, T cell activation altered a higher proportion of expressed genes, with 4,894 (33.4%) and 4,727 (32.3%) being upregulated and 4,931 (33.7%) and 4,611 (31.5) downregulated in CD4^+^ and CD8^+^ T cells, respectively, relative to the resting state (Fig. 2A). Interestingly, despite differences in their effector functions, CD4^+^ and CD8^+^ T cells shared a large proportion of activation-induced chromatin accessibility (Fig. 2B). Using annotations from the UCSC database in the TxDb.Hsapiens.UCSC.hg38.knownGene package (v3.4.6) and the ChromHMM model in the Roadmap Epigenomics Project ^[19]^, we annotated the activation-induced accessibility changes and calculated their enrichment over the universe of ATAC-seq peaks using GAT ^[20]^. Overall, approximately 75 % of the activation-induced peaks were located in intronic and intergenic regions and were significantly depleted for promoter regions (FDR < 0.05). Likewise, activation-induced peaks were enriched for ChromHMM predicted enhancer and other regulatory elements and were depleted in heterochromatin repressive marks and promoter-TSS regions compared to the universe of ATAC-seq peaks (Fig. S2A).

**Figure 2.**
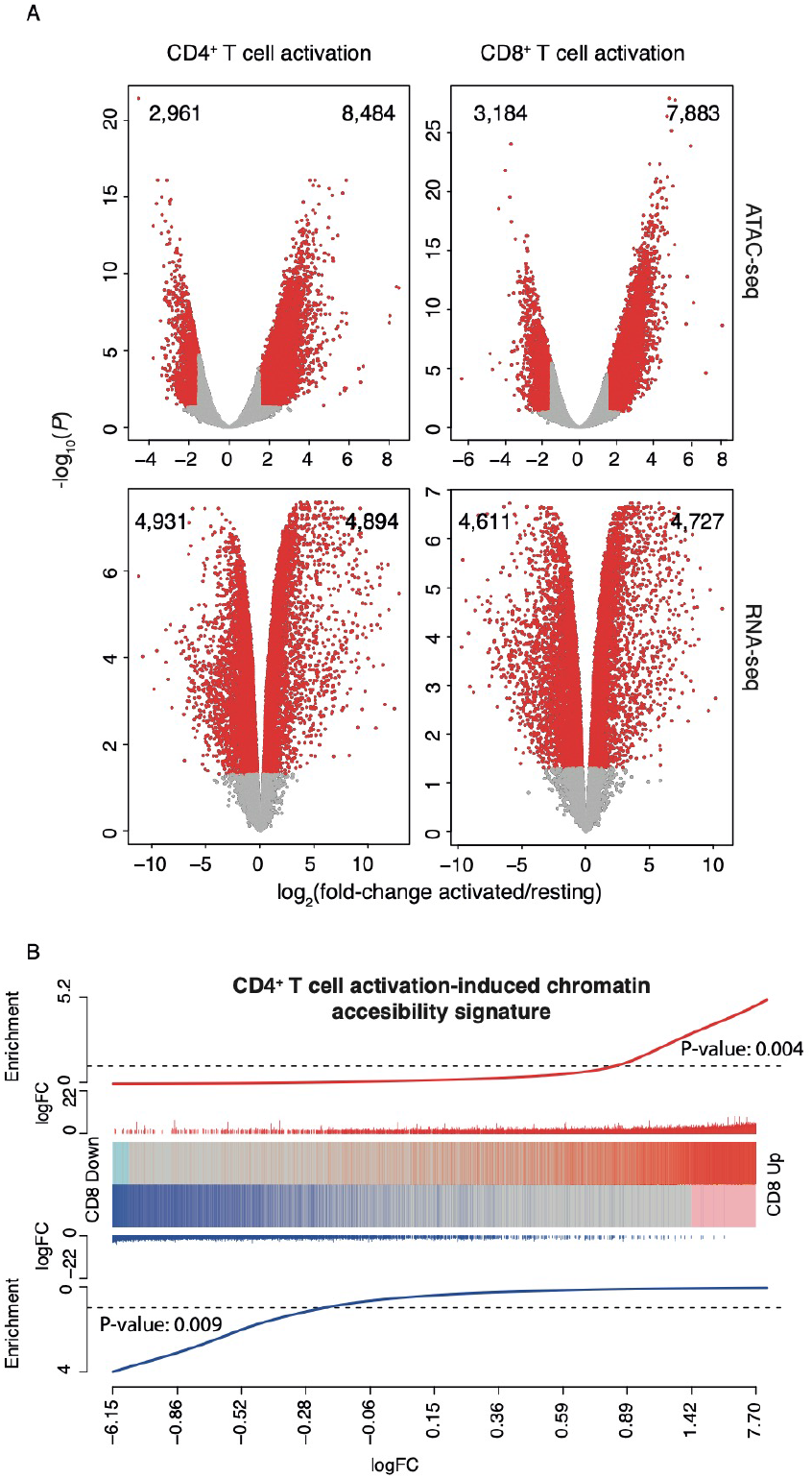
T cell activation induces genome-wide changes in chromatin and gene expression that are comparable between CD4^+^ and CD8^+^ T cells. (A) Volcano plots showing the effect of T cell activation for ATAC-seq (top) and RNA-seq (bottom), where the X-axis represents the log2 fold-change and the Y-axis the significance (−log10 P-value based on moderated t-statistics) for the activated and resting comparisons. Differentially accessible regions (FDR <0.05 and abs(logFC) ≥1.5) and differentially expressed genes (FDR < 0.05) are coloured in red. (B) Barcode enrichment plot showing strong correlation of the CD4^+^ T activation-induced accessibility signature in the CD8^+^ T cell activation-induced accessibility. Regions are ordered on the plot from right to left from most positive logFC to most negative upon CD8^+^ T activation (x axis). Vertical bars indicate regions significantly up (red) or down (blue) upon CD4^+^ cell T activation. Significant enrichment was assessed by fry gene set tests.

Several studies have shown that disease-associated SNPs frequently reside in cell-specific regulatory elements^[21–23]^. We therefore tested whether regions of altered chromatin accessibility associated with T cell activation were enriched for disease-/trait-associated SNPs or those in linkage disequilibrium (LD) with them (Table S2). After categorizing SNPs for 15 disease or trait classes we observed a strong enrichment for immune system, neurological autoimmune, haematological and cancer SNPs in activation-responsive elements compared to non-responsive elements (Fig. S2B). Moreover, enrichment was more pronounced in the subset of peaks that contained predicted enhancers by chromHMM (Fig. S2B).

To gain further insight into the activation-responsive chromatin elements in human T cells, we analysed activation-specific peaks for enrichment of known transcription factor binding (TFB) sites. Activation-induced peaks for both CD4^+^ and CD8^+^ T cells were specifically enriched for DNA motifs recognized by TFs involved in T cell development, activation or proliferation ^[10, 24–28]^. These included members of the bZIP (BATF, FOSL1, ATF3), NFKB (NFKB-p50, NFKB-p65), Zf (EGR2, EGR1), STAT (Stat5, Stat1, Stat3) and IRF (IRF4) families (Fig. S2C). Most changes in TFB site accessibility were accompanied by increased expression of the relevant TFs including those of the bZIP family, further supporting the regulatory nature of these activation-induced elements.

As whole transcriptome sequencing without poly(A) pre-selection allowed us to detect enhancer RNAs^[15]^, we could identify the subset of eRNAs that overlapped the set of human enhancer regions defined by the FANTOM consortium ^[29]^. In order to investigate if variation in chromatin accessibility was related to changes in eRNA expression, and thus enhancer activity, we calculated the distribution of eRNA expression across the activation-induced peaks. We observed that eRNA expression was on average significantly higher in the chromatin regions that gained accessibility upon activation than in those that lost accessibility (Fig. S3A).

To further explore the relationship between the activation-induced accessibility changes and expression of their potential target genes we applied the Genomic Regions Enrichment of Annotations Tool (GREAT) ^[30]^. Using GREAT we associated our set of activation-induced peaks with their putative target genes and observed that expression of genes linked to the peaks that gained accessibility was on average significantly higher than that of genes linked to peaks that lost accessibility (Fig. S3B). The set of target genes defined by GREAT comprised immune system-associated genes known to be upregulated upon T cell activation, including IL23R, IL12RB2, IL1R2, TNFRSF8, IL2, IL21 and IL13 among others. Furthermore, functional enrichment analysis using GREAT showed that the activation-induced peaks were significantly enriched for a number of relevant Gene Ontology terms, including regulation of immune system process, regulation of response to stimulus or regulation of cellular processes, among others.

In summary, this integrated analysis revealed that T cell activation leads to extensive remodelling of chromatin accessibility and the formation of active regulatory chromatin regions associated with TFs relevant to T cell biology, enhancer activity and expression of genes critical in T cell responses.

### T cell activation induces extensive changes in chromatin interactions

Chromatin interactions bring distal regulatory elements and genes into spatial proximity to regulate transcription. To determine differential chromatin interactions between samples we used the diffHic pipeline ^[31]^ which was designed for the detection of differential interactions from Hi-C data. It partitions the genome into bins and counts the number of read pairs mapping to each pair of bins. Each bin pair is then tested for significant changes in intensity between libraries, using the methods in the edgeR package ^[32]^. Upon activation, in CD4^+^ and CD8^+^ T cells, respectively, at 25kb resolution, we identified 17,965 and 15,496 differential chromatin interactions (DIs) with a significant increase in interaction counts which we refer to as ‘gain’ and 12,434 and 11,376 DIs with a significant decrease in interaction counts that we refer to as ‘loss’ (Tables S3A and S3B). Then to examine the relationship between genomic accessibility and interactions we selected DIs containing at least one activation-induced regulatory element in one anchor and a promoter of an expressed gene in the other. We found that accessibility was significantly increased for ‘gain’ interactions compared to ‘loss’ interactions following activation of either CD4^+^ and CD8^+^ T cell (Fig. 3A). Furthermore, when we examined if ‘gain’ or ‘loss’ chromatin interactions were associated with gene up-or down-regulation we found that gain interactions were linked to a higher percentage of genes upregulated while ‘loss’ interactions were linked to a higher percentage or genes downregulated (Chi-square: 35.0, p<0.05) (Fig. 3B). Thus, ‘gain’ and ‘loss’ interactions are linked with upregulation and downregulation, respectively, of corresponding target genes.

**Figure 3.**
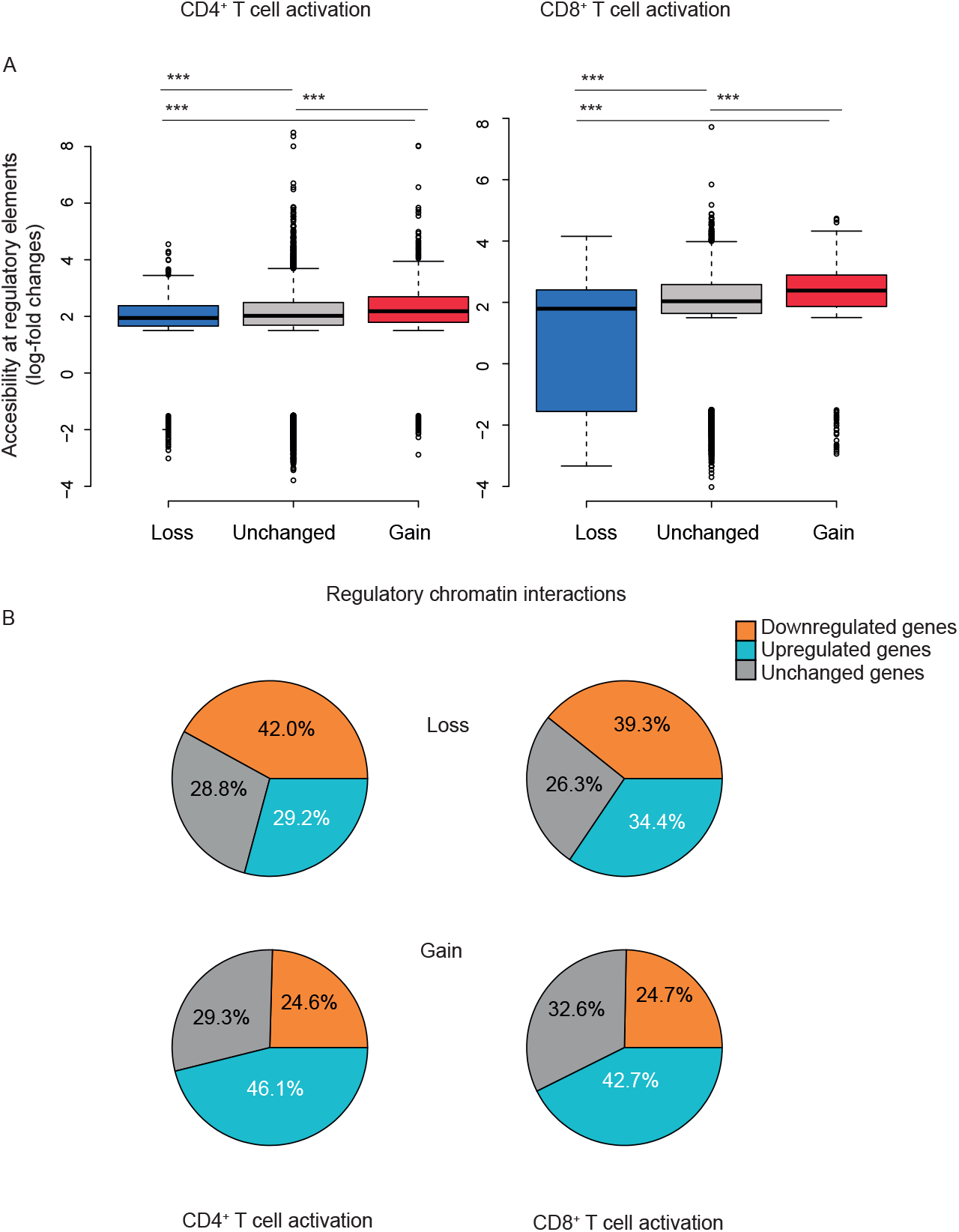
T cell activation induces extensive changes in gene-regulatory interaction. (A) Box plots showing chromatin accessibility at regions of differential interaction – ‘loss’, ‘unchanged’ or ‘gain’ - following T cell activation, at 25 kbp resolution (FDR<0.05). Shown are the median (central horizontal line), interquartile range (boxes), values of the upper and lower quartiles (whiskers), outliers beyond 1.5 IQR (circles). Statistical comparisons were made with the unpaired Wilcoxon test: ***P-value < 0.0001. (B) Pie chart showing distribution of downregulated, upregulated or unchanged target genes linked to ‘loss’ (top) or ‘gain’ (bottom) of regulatory interactions, for CD4^+^ (left) and CD8^+^ (right) T cells.

### T cell activation is associated with partitioning of genome topology

Evidence has emerged of the dynamic nature of TAD and DNA loop formation, mainly from studies of mouse embryo development (reviewed in ^[1]^). To examine TAD plasticity we first segmented the chromatin of naive and activated T cells, and B cells, into TADs using TADbit ^[33]^. Naïve CD4^+^ T, CD8^+^ T and B cells had 1,727, 1,750 and 1,602 TADs with a mean size of 1.40, 1.38 and 1.51 Mb, respectively, whereas activated CD4^+^ and CD8^+^ T cells had 2,772 and 2,863 TADs with a mean size of 0.85 and 0.84 Mb, respectively (Fig. 4A). In activated T cells, TADs appeared to be partitioned, becoming smaller and more numerous compared to naïve T cells (TAD sizes: t = 11.3 and 10.7 for nCD4 *vs*. aCD4 and nCD8 *vs*. aCD8, respectively; P-value < 2e-16) (Fig. 4A). On average 62% of the TADs called in resting CD4^+^ T cells intersected with TADs in resting CD8^+^ T cells, with >75% reciprocal region overlap. Interestingly, this fell to 43.5 % when comparing the TAD overlap between resting and activated T cells. Further analysis of the TAD intersects revealed that while only 24% of the TADs called in resting CD4^+^ T cells overlapped two or more TADs in resting CD8^+^ T cells this increased to 44% when analysing the TAD overlap between resting and activated T cells (Fig. S4), thus suggesting that T cell-activation induces partitioning of TADs into finer chromatin domains.

**Figure 4.**
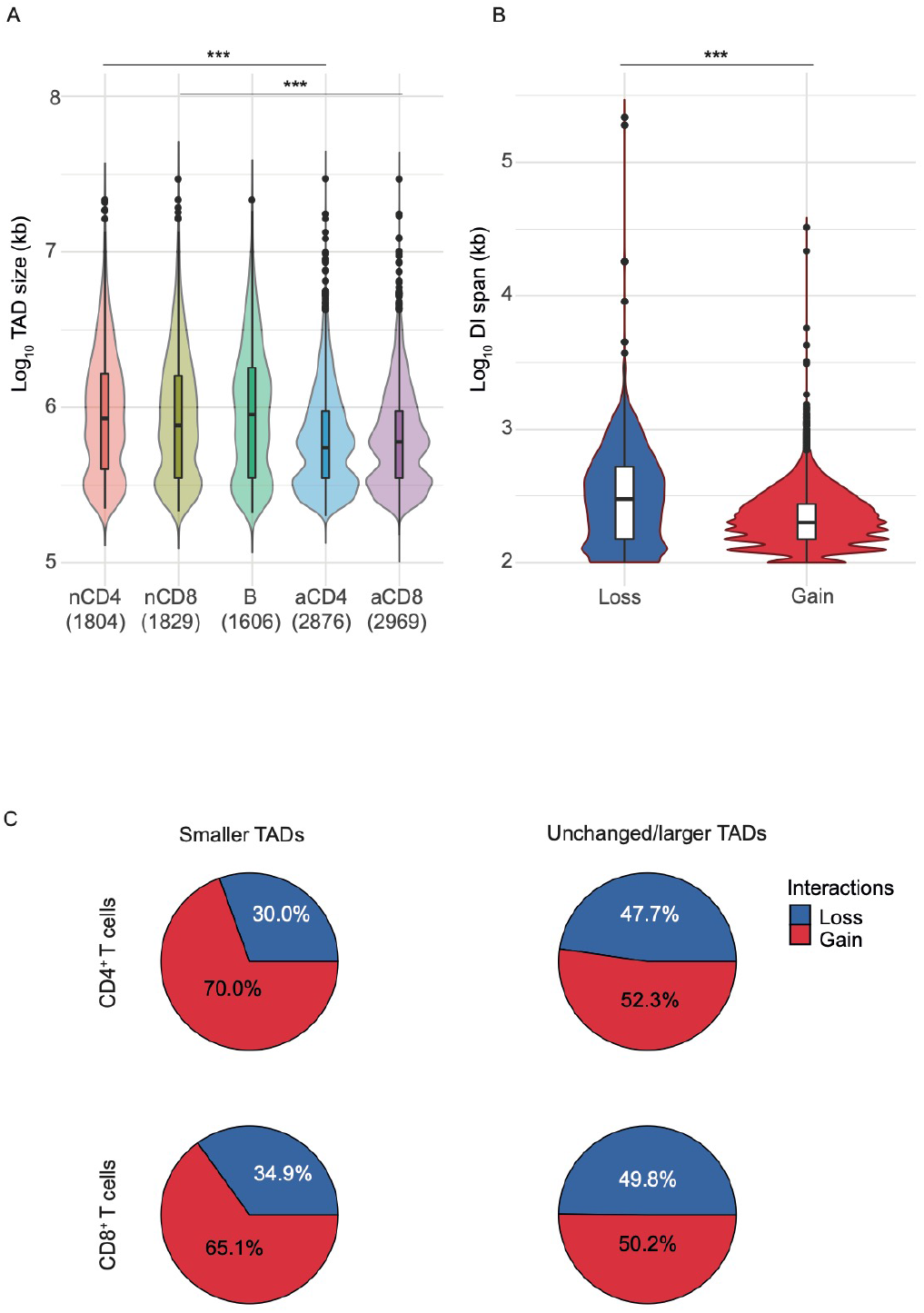
T cell activation results in partitioning of genome topology. (A) Violin plots showing the distribution of TAD sizes in resting and activated T cells and B cells determined from summed libraries called with TADbit. Numbers in parenthesis depict the number of TADs called in each category. (B) Violin plots showing the distribution of the span of differential chromatin interactions (DIs) determined at 25 kbp resolution (FDR<0.05) that are lost (decrease in logFC) or gained (increase in logFC) upon activation. Violin plots show median (horizontal line), interquartile range (boxes), values of the upper and lower quartiles (whiskers), outliers (circles) and kernel density estimation. Data for CD4^+^ and CD8^+^ T cells were merged as they showed a similar pattern. (C) Pie chart showing distribution of Loss and Gain chromatin interactions in TADs that become smaller in size (left) or remained unchanged/became larger in size (right) upon T cell activation. Statistical comparisons were made with the unpaired Wilcoxon test: ***P-value < 0.0001

Furthermore, we also observed that, analogous to TAD partitioning, chromatin interactions strengthened upon activation were on average significantly shorter than those that were weakened (t= 9.3; P-value < 2e-16), i.e. activation preferentially promoted disruption of the longer-range interactions (Fig. 4B). Remarkably, intra-TAD chromatin interactions in TADs that became smaller upon activation showed a higher proportion of ‘gain’ interactions than those in TADs that remained unchanged or larger (Fig. 4C), thus suggesting that chromosome domains play a pivotal role in shaping the intra-TAD chromatin loops. Moreover, TAD partitioning and rearrangement of chromatin interactions overlapped markers of T cell activation including CDH3, CDH1, IL23R and IL12RB2 (Fig. 5, Fig. S5).

**Figure 5.**
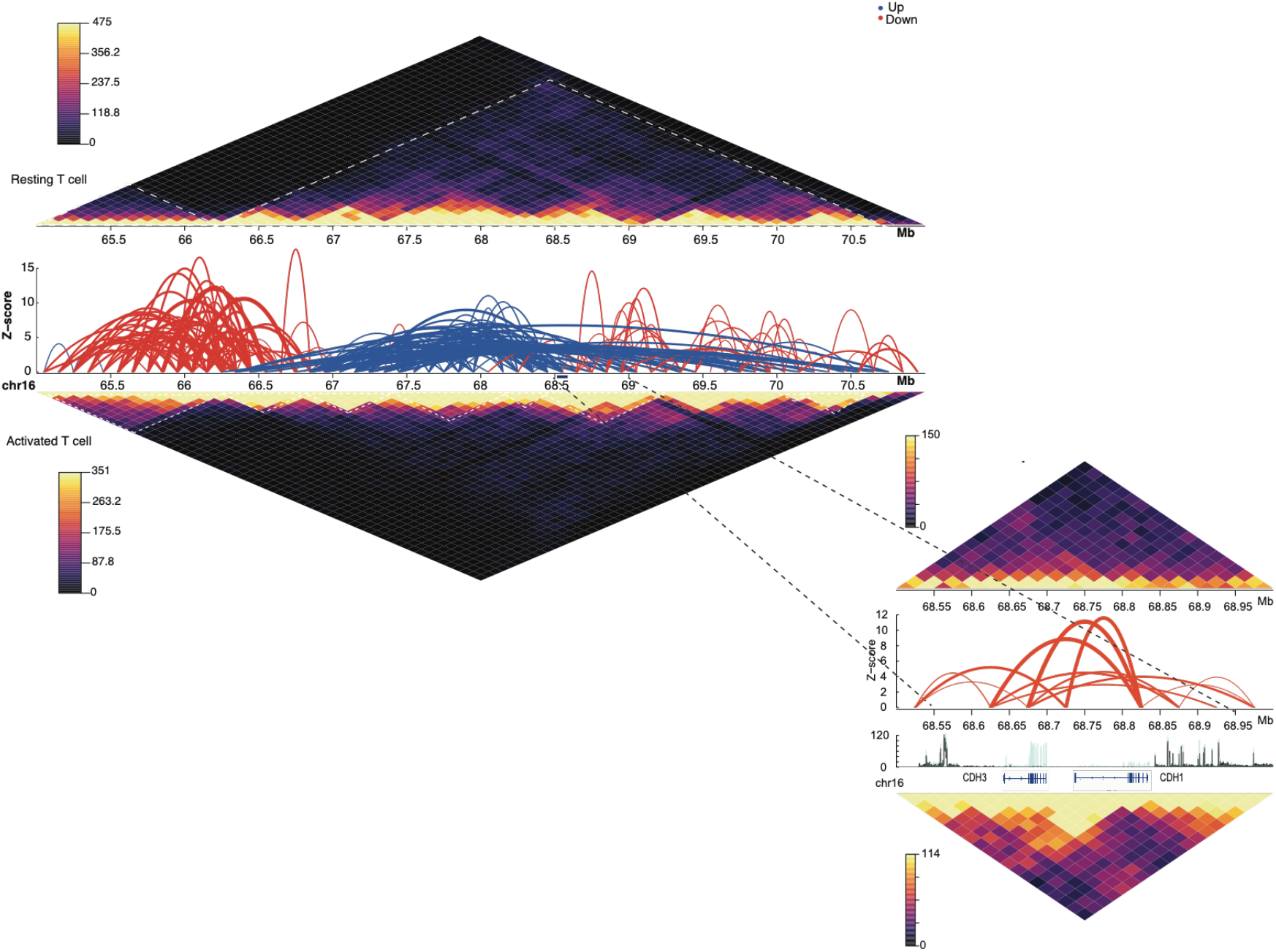
T cell activation results in partitioning of genome topology. *In-situ* Hi-C contact matrices plotted at 100 kbp resolution in resting and activated T cells over genes that have been associated with T cell activation such as CDH3 and CDH1. Colour scale indicates number of reads per bin pair. Locations of the TADs as called by TADbit at these regions are indicated below each contact matrix. Significant differential interactions at 100 kbp (FDR<0.05) are represented by arches where the z-score is −log10(p-value). Red and blue lines represent gain and loss of interactions, respectively, in response to activation. The zoomed inset regions show in-situ Hi-C contact matrices, significant differential interactions at 25 kbp resolution (FDR<0.05) and RNA sequencing coverage plots at the CDH3 and CDH1 loci. Turquoise and black coverage plots represent activated and resting T cells, respectively.

We next sought to identify features associated with the strengthening and weakening of TAD boundaries. We defined differential TAD boundaries, using the diffHic package ^[31]^. To identify these boundaries, we use the directionality index defined by Dixon et al.^[5]^. For each region the directionality index was computed by counting the number of read pairs mapped between the target locus and an ‘upstream’ interval with higher genomic coordinates; counting the number of read pairs mapped between the target locus and a ‘downstream’ interval with lower genomic coordinates; and taking the normalized difference of the counts. We then used the statistical framework of edgeR to identify regions (boundaries) where upon activation the size or direction of the directionality statistic changes. Upon activation, 5,278 and 4,705 boundaries became significantly stronger and 6,600 and 7,312 weaker in CD4^+^ and CD8^+^ T cells, respectively. We then calculated the distribution of chromatin accessibility, nucleosome occupancy and gene expression for regions containing differential TAD boundaries in both resting and activated T cells. Interestingly, we observed that TAD boundaries that became significantly stronger upon activation were associated on average with lower chromatin accessibility, higher nucleosome occupancy and lower protein-coding gene expression (Fig. 6). In summary, T cell activation leads to global rearrangement of TAD boundaries, partitioning of TADs and disruption of long-range chromatin interactions, with strengthening of TAD boundaries associated with lower chromatin accessibility, higher nucleosome occupancy and lower gene expression.

**Figure 6.**
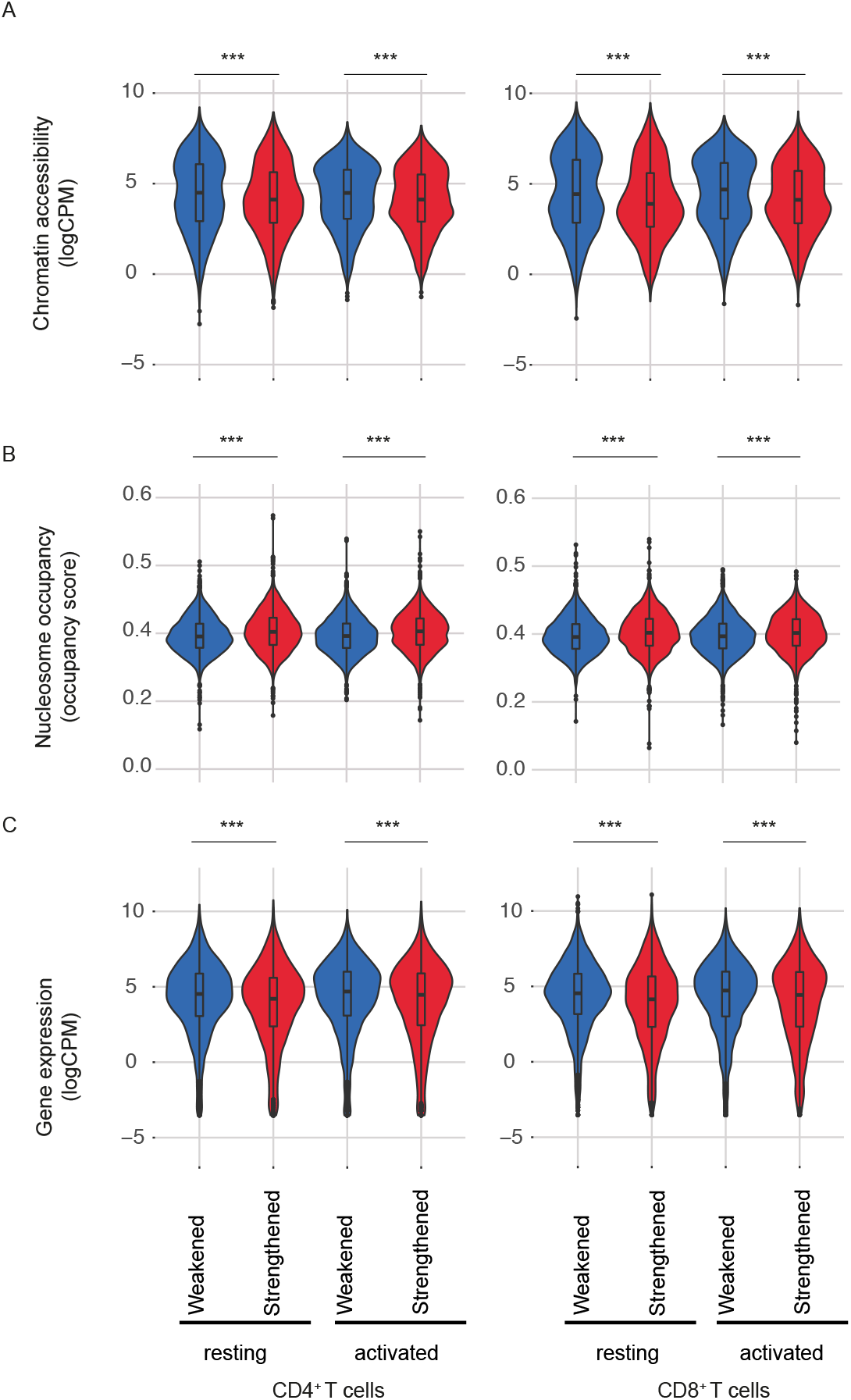
Increased strength in TAD boundary and chromatin interactions upon T cell activation is associated with nucleosome occupancy and gene expression. Violin plots showing the distribution of (A) chromatin accessibility (logCPM), (B) nucleosome occupancy (NucleoATAC occupancy score) and (C) expression of protein-coding genes (logCPM) across differential TAD boundaries strengthened and weakened following activation of CD4^+^ and CD8^+^ T cells. Violin plots show median (horizontal line), interquartile range (boxes), values of the upper and lower quartiles (whiskers), outliers (circles) and kernel density estimation. Statistical comparisons were made with the unpaired Wilcoxon test: ***P-value < 0.0001.

### Chromosome A/B compartments are not altered substantially by T cell activation

Lineage differentiation has been associated with rearrangement of chromosome A/B compartments ^[34]^. However, in response to T cell activation we observed only minor changes in the A/B compartments, with less than 4% of the genome switching compartments (Fig. 7A, 7B). As expected, the small proportion of regions that switched from A to B showed decreased accessibility and gene expression, whereas regions that switched from B to A tended to show higher accessibility and gene expression (Figure 7C). Changes across T and B cell types were also minimal. Overall, the pattern of change was subtle, indicating that T cell activation has minimal impact on the A and B compartments.

**Figure 7.**
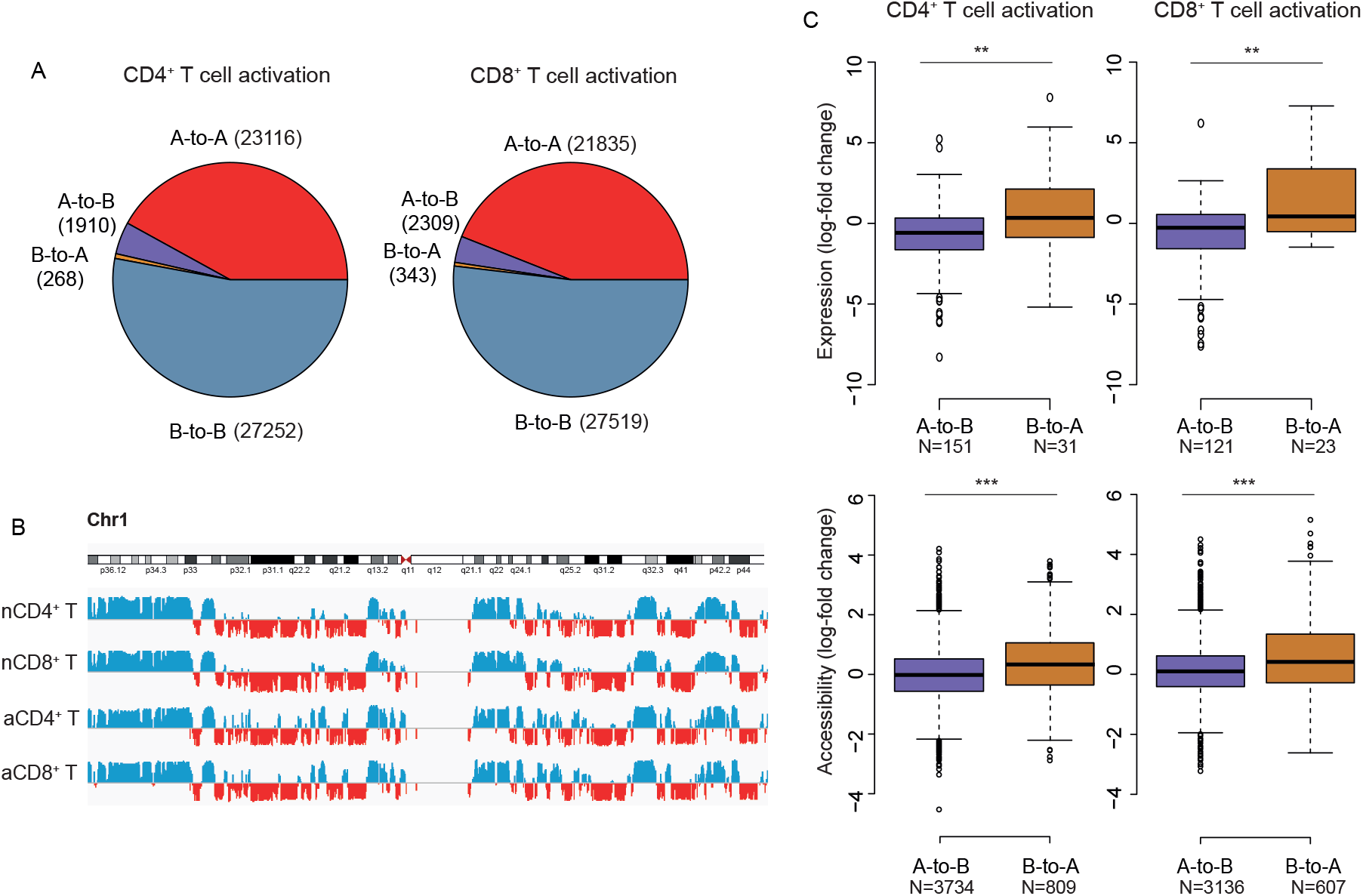
Chromosome A/B compartments are not altered substantially by T cell activation. (A) Pie charts showing the distribution of chromatin regions in the four different group of compartment changes: stable (A to A, B to B) or flipping (A to B and B to A). This was defined by pairwise comparison between activated and naive T cells. (B) PCA (principal component analysis) scores in resting and activated T cells along chromosome 1 representing the stability of the A/B compartments. PCA scores were calculated as described in Methods. (C) Boxplots showing the distribution of chromatin accessibility and gene expression at regions that exhibit A/B compartment shifts upon T cell activation. Shown are the median (central horizontal line), interquartile range (boxes), values of the upper and lower quartiles (whiskers), outliers beyond 1.5 IQR (circles). Statistics were determined with the unpaired Wilcoxon test. ***P-value < 0.0001.

## DISCUSSION

In this study, we analysed chromatin accessibility, Hi-C chromatin interactions and the whole transcriptome in human CD4^+^, CD8^+^ T cells under resting (naïve) and activated conditions, and in B cells. Our study made use of biological replicates throughout, so that changes in chromatin accessibility, interaction intensity or gene expression were assessed using statistically rigorous methods relative to the biological variation observed between human individuals. Activation of T cells was associated with multi-level remodelling of chromatin structure, that was comparable between CD4^+^ and CD8^+^ T cells. It entailed extensive changes in chromosome accessibility in regions enriched for regulatory elements and eRNAs, immune system-associated SNPs and TFBSs recognized by TFs involved in T cell development, activation or proliferation. What we demonstrate in human T cells parallels what has been described in other systems ^[12–14,35,36]^, namely that changes in chromatin accessibility in response to external cues identify active regulatory regions likely to be critical for transcriptional responses. Furthermore, we show that a T cell activation uncovers many additional regulatory regions likely to be associated with the immune response. T cell activation also induced thousands of differential chromatin interactions that were linked to the upregulation or down regulation of the corresponding target genes. At a higher level of chromosome architecture, T cell activation was linked with genome-wide rearrangement of TAD boundaries. Previous studies have either characterized activation-responsive chromatin accessibility ^[13,14,35,36]^ or 3D organization of the genome ^[8,13,15]^ in mouse and human immune cells. However, to the best of our knowledge, no study has previously integrated chromatin accessibility, nucleosome occupancy and 3D genome structure in relation to gene transcription in response to activation of human T cells. Ours is also the first study to investigate activation-induced changes in chromatin architecture in the two main subpopulations of T cells, CD4^+^ and CD8^+^ T cells, enabling us to determine if their activation is linked to similar genome control mechanisms.

Although TADs were reported to be largely invariant across species and cell types ^[5,6]^, recent evidence ^[1]^ indicates they are more dynamic. In fact, we found that T cell activation was associated with global rearrangement of chromatin interactions involving changes in the strength of TAD boundaries and a finer compartment structure as indicated by the smaller size of TAD domains. Strengthened TAD boundaries displayed lower chromatin accessibility, higher nucleosome occupancy and lower gene expression both before and after activation. Nucleosome organization is known to dictate local chromatin folding by regulating internal factors including linker DNA length and linker-histone binding affinities, but whether nucleosome architecture influences higher level genome topology, i.e. TADs and DNA loops, remains unknown ^[37]^. Polymer simulations of the chromatin fiber suggest that regions of “stiffness” can act as local insulators to decrease interaction frequencies ^[38]^. In mammary epithelial MCF-10A cells, knockdown of BRG1, the major ATPase subunit of the SWI/SNF chromatin remodelling complex, globally altered long-range genomic interactions and decreased TAD boundary strength, suggesting that nucleosome occupancy around CTCF sites might contribute to the regulation of higher-order chromatin architecture ^[39]^. Our results in primary human cells support the view that long-range chromosomal organization and nucleosome occupancy are molecularly coupled and that regions of TAD boundary formation are primed before activation as indicated by the lower accessibility and higher nucleosome occupancy and in both resting and activated T cells. Further examination of other elements at differential TAD boundaries may provide valuable insights into the TAD formation process and its role in chromatin folding and transcriptional regulation.

Together, our findings connect features of chromatin structure to gene expression in response to T cell activation. In addition, we provide a genome-wide resource of T cell activation-induced regulatory regions that will aid interpretation of chromatin accessibility data in human T cells.

## ACKNOWLEDGEMENTS

We thank Dr. Stephen Wilcox (Genomics Hub, WEHI) for generating the sequencing data and Drs. Natasha Jansz and Marnie Blewitt for their advice on the ATAC-seq protocol. This work was supported by Australian National Health and Medical Research Council (NHMRC) Program Grant (APP1037321), Research Fellowships (H.D.C, T.M.J, L.F, G.K.S, R.S.A, L.C.H) and Centre for Research Excellence Grant (APP 1078106) (N.G.B.). The work was made possible through Victorian State Government Operational Infrastructure Support and NHMRC Research Institute Infrastructure Support Scheme.

## AUTHOR CONTRIBUTION

L.C.H and N.G.B designed the project and wrote the manuscript. G.N. and T.M.J performed the RNA-seq, ATAC-seq and Hi-C experiments. N.G.B, H.D.C, A.L.G, J.S. performed the computational analyses. G.K.S, R.S.A, E.B.S and L.F. contributed to data interpretation and manuscript editing.

## METHODS

### Blood collection, cell subset isolation and T cell activation

Heparinised venous blood was collected at 09.00 h from two healthy young adult males, who gave written informed consent. The research was carried out in accordance with the principles of the Declaration of Helsinki and the guidelines of Nature journals and was approved by the Walter and Eliza Hall Institute of Medical Research Human Research Ethics Committee (application 88/03). Peripheral blood mononuclear cells (PBMCs) were purified by Ficoll-Hypaque gradient centrifugation and cryopreserved in liquid N2. Thawed PBMCs, ≥92% viable by acridine orange-ethidium bromide staining, were stained with anti-human αβ TCR (eBioscience, clone IP26, cat. no. 46-9986-42), anti-human CD4 (BD Pharmingen, clone RPA-T4, cat. no. 555349), anti-human CD45RA (BD Pharmingen, clone 5H9, cat. no. 556626), anti-human CD25 (BD Pharmingen, clone M-A251, cat. no. 557741), anti-human CD14 (BioLegend, clone 63D3, cat. no. 367104), anti-human CD16 (BD Pharmingen, clone 3G8, cat. no. 557758), anti-human HLA-DR (eBioscience, clone L243, cat. no. 48-9952-42), and anti-human CD19 (BioLegend, clone HIB19, cat. no. 302238). Naïve CD4^+^ T cells (CD14^−^CD16^−^TCRαβ^+^CD4+^−^CD45RA^+^CD25^−^) and CD8^+^ T cells (CD14^−^CD16^−^TCRαβ^+^CD4+^−^CD45RA^+^CD25^−^), and B cells (TCRαβ^−^, HLA-DR^+,^ CD19^+^), were flow-sorted on a FACSAria (BD Biosciences). Naive CD4^+^ and CD8^+^ T cells were cultured in Iscove’s Modified Dulbecco’s Medium containing 5% pooled, heat-inactivated, human serum, 100 nM non-essential aminoacids, 2mM of glutamine and 50 uM 2-mercaptoethanol (IP5) medium. T cells were activated through the T cell receptor subunit CD3 and the co-stimulator CD28 by the addition of Human T-Activator CD3/CD28 Dynabeads at a 1:2 bead:cell ratio (Life Technologies, cat. no. 111.31D) for 72 hours.

### RNA-seq

RNA was isolated using the miRNeasy Micro Kit (QIAGEN, cat. no. 217084). RNA libraries were prepared with an Illumina's TruSeq Total Stranded RNA kit with Ribo-zero Gold (Cat# RS-122-2001, Illumina) according to the manufacturer’s protocol. rRNA-depleted RNA was purified and then reverse transcribed into cDNA using SuperScript II reverse transcriptase (Cat# 18064014, Invitrogen). Validation of the library preparations was performed using Agilent D1000 ScreenTape Assay on the Agilent 2200 TapeStation. Total RNA-Seq libraries were sequenced on the Illumina NextSeq 500 platform to produce 2 × 75 paired-end reads. All samples were aligned to the human genome, build hg38, using the Rsubead aligner v1.24.1 ^[40]^. The number of fragments overlapping each Entrez gene were summarised using featureCounts ^[41]^ and Rsubread's inbuilt hg38 annotation. Gene symbols were associated with Gene IDs using NCBI gene information (ftp://ftp.ncbi.nlm.nih.gov/gene/DATA/GENE_INFO).

### ATAC-seq

ATAC-seq was performed as previously described ^[16]^. For each immune cell subset, 50,000 purified cells were lysed in cold lysis buffer (10 mM Tris-HCl, pH 7.4, 10 mM NaCl, 3 mM MgCl_2_ and 0.03% Tween20). Immediately after lysis, nuclei were spun out at 500 g for 8 minutes at 4°C and the supernatant carefully removed. Nuclei were resuspended with Tn5 transposase reaction mix (25 μl 2X TD buffer, 2.5 μl Tn5 transposase, and 22.5 μl nuclease-free water) (Nextera DNA Library Prep Kit (Illumina, cat. no. FC-121-1030). The transposase reaction was performed at 37°C for 30 minutes, and DNA immediately purified with a Qiagen MinElute kit (QIAGEN, cat. no. 28204). ATAC-seq libraries were sequenced on an Illumina NextSeq 500 to produce 75bp paired-end reads.

### In situ Hi-C

In situ HiC was performed as previously published ^[42]^. Primary immune cell libraries were generated in biological duplicates. Libraries were sequenced on an Illumina NextSeq 500 to produce 81bp paired end reads. Between 129 million and 280 million valid read pairs were generated per sample.

### Differential gene expression analysis

Differential expression analyses were undertaken using the edgeR v3.20.9 ^[32]^ and limma v3.34.9 ^[43]^ software packages. Any gene which did not achieve a count per million mapped reads (CPM) greater than 1.5 in at least 2 samples were deemed to be unexpressed and subsequently filtered from the analysis. Additionally, all genes without current annotation were also removed. Compositional differences between libraries were normalized using the trimmed mean of log expression ratios method (TMM)^[44]^. All counts were then transformed to log2-CPM with associated precision weights using voom ^[45]^. Differential expression between all cell types was assessed using linear models and robust empirical Bayes moderated t-statistics ^[46]^. To increase precision, the linear models incorporated a correction for a donor batch effect. P-values were adjusted to control the FDR below 5% using the Benjamini and Hochberg method.

### eRNA analysis

Reads overlapping the non-exonic ATAC-seq peaks were summarised using featureCounts in the naive and activated CD4 and CD8 samples. To avoid inflation of expression estimates, the library size for each sample was set to the total number of reads aligned to the genome for that sample. All regions that failed to achieve a CPM greater than 0.5 in at least 2 samples were considered to be unexpressed and were therefore filtered from the analysis. The TMM method was then applied to normalize compositional differences between libraries and the data transformed to log2-CPM with precision weights using voom. Differential expression of the regions was then evaluated between the activated and naïve CD4^+^ and CD8^+^ T cell samples using linear models and robust empirical Bayes moderated t-statistics. P-values were adjusted to control the FDR below 5% using the Benjamini and Hochberg method.

### ATAC-seq data pre-processing and peak calling

75 bp ATAC-seq reads were aligned to the human genome assembly (hg38) using Bowti2 v2.2.5 (bowtie2 -p 4 -X 2000) ^[47]^. For each sample, mitochondrial reads, ummapped reads and low mappability (<30) reads were filtered out using Samtools (v1.6) function “view” ^[48]^. After filtering, we had a median of 80 million (MAD +/− 13 million) reads per sample. Filtered ATAC-seq reads from naive and activated CD4^+^ and CD8^+^ T cell as well as B cells from two donors were merged using samtools function merge, and peaks were called on the merged bam file using MACS2 (v2.1.0) –callpeak (with parameters -- nomodel, --extsize 200, and --shift 100) ^[49]^, such that there were 113,689 peaks after excluding peaks mapping outside the main chromosome contigs. ATAC-seq reads overlapping the peaks were summarized using featureCounts ^[41]^. Peaks in blacklisted genomic regions as defined by ENCODE for hg38 were removed.

### Differential accessibility analysis

Differential accessibility analysis was undertaken using the edgeR v3.20.9 ^[50]^ and limma v3.34.9 ^[43]^ software packages. The TMM method was applied to normalize compositional differences between libraries ^[44]^. A mean-dependent trend was fitted to the negative binomial dispersions with the estimateDisp function and, and differential accessibility between all cell types was assessed using likelihood ratio tests (LRT) in edgeR. As is the differential expression analysis, linear models incorporated a correction for a donor batch effect. P-values were adjusted for multiple testing using the Benjamini-Hochberg method. Peaks with a FDR below 5% and abs(logFC) ≥1.5 were defined as differentially accessible regions. Concordance between activation profiles was assessed by gene set tests using fry function in edgeR^[51]^.

### Enrichment of transcription factor binding motifs

The Homer suite v4.10 ^[52]^ was used to determine transcription factor enrichment within ATAC peaks, using the findMotifsGenome.pl function (with parameters hg38 and –size given).

### Annotation of ATAC peaks

Peaks were annotated as 5’, 3’, promoter-TSS, exonic, intronic, TTS or intergenic using the Homer suite annotatePeaks.pl function and the default setting. We also found all instances of TFBSs with altered accessibility upon activation in the ATAC-seq peaks using the annotatePeaks.pl function, and options –m. Chromatin state(s) of the peaks were annotated using the ChromHMM calls for naive CD4^+^ and T CD8^+^ T cells and B cells from the Roadmap Epigenomics Project with BEDtools (function intersectBed) ^[53]^. Functionally-related subclasses were aggregated as follows: Promoter-TSS (TssA, PromU, PromD1 and PromD2), transcribed (Tx5', Tx, Tx3', TxWk), enhancers and other regulatory elements (TxReg, TxEnh5', TxEnh3', TxEnhW, EnhA1, EnhA2, EnhAF, EnhW1, EnhW2 and EnhAc), repetitive elements (ZNF/rep), heterochromatin/repressive (Het, ReprPC, Quies) and others (DNAse, PromP, PromBiv).

### Enrichment for chromatin states and gene annotation

Enrichment of ATAC-seq peaks for chromatin states as well as for gene annotations was calculated using GAT v1.3.4 ^[54]^. The significance threshold was set up at FDR below 5%.

### Prediction of transcriptional target genes

To identify potential targeted genes for the activation-induced peaks we performed enrichment analysis of gene annotations in the proximity of the activation-induced peaks using Genomic Regions Enrichment of Annotations Tool (GREAT) ^[30]^ against the whole genome as background. GREAT links genomic regions with genes by defining a ‘regulatory domain’ for each gene in the genome. Gene regulatory domains were defined with the “Basal plus extension” association rules (proximal 5 kb upstream and 1 kb downstream from the TSS, plus distal extended to the nearest gene’s basal domain but not more than 500kb). Significantly enriched gene sets were then selected by FDR < 0.05 for binomial tests to identify the regulatory domains with the densest clusters of activation-induced peaks and classify them as potential transcription target genes.

### Enrichment of GWAS loci

GREGOR v1.4.0 ^[55]^ was used for enrichment analysis of disease-trait associated SNPs in the ATAC-seq peaks. GREGOR calculates enrichment relative to MAF, TSS-distance and number of LD neighbor-matched null SNP sets using the GREGOR parameters: r^2^ threshold=0.8, LD window size=1 Mb and minimum neighbor number= 500. For the GWAS SNPs, we created an updated version of Table S2 in Maurano et al ^[22]^. The GWAS SNP set used for analysis was derived from the NHGRI GWAS Catalog, downloaded on August 2, 2017. The catalog contained 41,304 entries at the time of download. We excluded SNPs mapping outside the main chromosome contigs, including the “random” chromosome fragments, SNPs without coordinates in the GRCh37/hg19 human genome assembly. There were 40,929 unique SNP disease/trait combinations that represented 34,421 unique SNP IDs (Table S2). Of these, 19,075 were in non-coding regions. Coding regions were defined by the CCDS Project (downloaded from the UCSC genome browser at http://hgdownload.cse.ucsc.edu/goldenPath/hg38/database/ccdsGene.txt.gz on August 4, 2017). As in Maurano et al ^[22]^, we also grouped SNPs into classes of similar diseases or traits but some of the categories were updated so they could better reflect the new and extended list of SNPs. Categories comprised: aging related; immune system; cancer; cardiovascular diseases and traits; metabolic disorder; drug metabolism; hematological parameters; kidney, lung, or liver; miscellaneous; serum metabolites; neurological/behavioral; neurological/autoimmune; parasitic or bacterial disease; quantitative traits; radiographic (primarily bone density); viral disease.

### Nucleosome occupancy

Nucleosome dyad centers were identified in the “nucmap_combined.bed” file, while nucleosome occupancy was defined with the “occ.bedgraph” files using the NucleoATAC v0.3.2.1 package ^[56]^.

### In situ Hi-C data pre-processing

Each sample was aligned to the hg38 genome using the presplit_map.py script in the diffHic package v1.10.0 ^[31]^. Data were pre-processed and artefacts removed as per Johanson et al. 2018 ^[57]^.

### Differential interaction (DI) analysis

DIs between all five libraries were detected using the diffHic package ^[31]^ at two different resolutions, 100 kbp and 25 kbp. Read pairs were counted into 25 or 100 kbp bin pairs (with bin boundaries rounded up to the nearest MboI restriction site) using the squareCounts function. This yielded a matrix of read pair counts for each bin pair in each library. All bins with counts less than 5 were discarded along with bins on the sex chromosomes. Bins containing blacklisted genomic regions as defined by ENCODE for hg38 ^[58]^ were also removed. Filtering of bin-pairs was performed using the filterDirect function, where bin pairs were only retained if they had average interaction intensities more than 6-fold higher than the background ligation frequency. The ligation frequency was estimated from the inter-chromosomal bin pairs from a 2 Mbp bin-pair count matrix. Bins on the first diagonal of the interaction space are also removed with the filterDiag function.

For the retained bin pairs, counts were normalized between libraries using a LOESS-based approach to account for abundance-dependent biases. This was performed using the normOffsets function to obtain a matrix of offsets with bin pairs less than 100 kbp (for the 25kbp) or 150 kbp (for the 100 kbp) from the diagonal normalized separately from other bin pairs. Tests for differential interactions were performed using the quasi-likelihood (QL) framework^[59,60]^ of the edgeR package (v3.20.9), which assesses statistical significance relative to biological variation between the replicate libraries. The design matrix was constructed using a layout that specified the cell lineage to which each library belonged and which individual the cells were sampled from. Using the counts and offsets for all bin pairs, a mean-dependent trend was fitted to the negative binomial dispersions with the estimateDisp function. A generalized linear model (GLM) was fitted to the counts for each bin pair ^[61]^, and the QL dispersion was estimated from the GLM deviance with the glmQLFit function. The QL dispersions were then squeezed toward a second mean-dependent trend, using a robust empirical Bayes strategy ^[46]^ to share information between bin pairs. A P-value was computed for each bin pair using the QL F-test, representing the evidence against the null hypothesis, i.e., no difference in counts between groups. P-values were adjusted for multiple testing using the Benjamini-Hochberg method. If a bin pair had a FDR below 5%, it was defined as a DI. To reduce redundancy in the results, DIs adjacent in the interaction space were aggregated into clusters using the diClusters function to produce clustered Dis. DIs were merged into a cluster if they overlapped in the interaction space, to a maximum cluster size of 500 kbp (for the 25kbp) or 1Mbp (for the 100 kbp) to mitigate chaining effects. The significance threshold for each bin pair was defined such that the cluster-level FDR was controlled at 5%. Cluster statistics were computed using the combineTests and getBestTest functions from the csaw package v1.12.0 ^[62]^. Clustered DIs were used to report the number of differences between the libraries. The 25kbp unclustered DIs were used for overlap analysis and integration with other data types. Contact matrices were created from the libraries using the inflate function in diffHic for various bin sizes with no filtering. Contact matrices from biological replicates were summed.

### Detection of TADs

TAD were detected with the TADbit v0.2.0.5 python based software ^[33]^. Read pairs were counted into 50 kbp bin pairs (with bin boundaries rounded up to the nearest MboI restriction site) using the squareCounts function of diffHic with no filter. This yielded a count matrix containing a read pair count for each bin pair in each library. The count matrix was converted into a contact matrix for each somatic chromosome with the inflate function of the InteractionSet package ^[62]^. Replicate contact matrices were summed. TADs were detected for each chromosome with the function find tad of the TADbit software. Only TADs boundaries with a score of 7 or higher were included in the results.

### Detection of differential TAD boundaries

Differential TAD boundaries between all five libraries were detected using the diffHic package ^[31]^. To identify boundaries, the directionality index ^[5]^ for each genomic region was determined by counting the number of read pairs mapped between the genomic region 1 Mbp upstream and separately 1Mbp downstream then taking the normalized difference of the counts. This produces a log-fold-change for each sample at each genomic region. Differential TAD boundaries were determined to be genomic regions were the size or the direction of the logFC between up and down stream counts changed significantly between conditions. The domainDirections function was used to determine counts for 100 kbp genomic regions for 1 Mbp up- and downstream. A DGEList was constructed from the counts and library size. Low average abundance regions (calculated across all samples) of less than 0.8 were removed. Tests for differential TAD boundaries were performed again using the QL framework of the edgeR package as described in the differential interaction analysis section. A design matrix was constructed with cell type specific coefficients for the log-fold change between up- and downstream counts for each cell type. It also contains library specific and individual blocking factors. A P-value was computed for each 100kbp region using the QL F-test and adjusted for multiple testing using the Benjamini-Hochberg method (FDR below 5%). A differential TAD boundary was considered to be strengthened from one cell type to another if the absolute value of the log-fold change between up- and downstream has increased.

### Detection of A/B compartments

A/B compartments were identified at a resolution of 50 kbp using the method outlined by Lieberman-Aiden et al. ^[63]^ using the HOMER HiC pipeline ^[52]^.

After processing with the diffHic pipeline libraries were converted to the HiC summary format using R. Then input tag directions were created for each library with the makeTagDirectory function specifying the genome (hg38) and restriction enzyme cute site (GATC). Biological replicates tag directories for each cell type were summed. The runHiCpca.pl function was used on each library with -res 50000 and specifying the genome (hg38) to perform PCA to identify compartments. To identify changes in A/B compartments between libraries, the getHiCcorrDiff.pl function was used to directly calculate the difference in correlation profiles. We identified regions with an A-to-B or B-to-A compartment flip that showed a correlation profile lower than 0.6 and regarded them as differential compartments.

### Visualization of the data and plots

Plots were performed using R and ggplot. Multi-dimensional scaling (MDS) plots were constructed using the plotMDS function in the limma v3.34.9 package ^[43]^ applied to the filtered and normalized log2-counts-per-million values for each library. The removeBatchEffect function of the limma package was used to correct for effect of the individual and batch in the data. Plaid plots were constructed using the contact matrices and the plotHic function from the Sushi v1.22.0 R package ^[64]^. The inferno color palette from the viridis v0.5.1 package ^[65]^ was used and the range of color intensities in each plot was scaled according to the library size of the sample, to facilitate comparisons between plots from different samples. DI arcs were plotted with the plotBedpe function of the Sushi package. The z-score shown on the vertical access was calculated as −log10 (P-value). RNA-seq coverage was plotted with the plotBedgraph function of the Sushi package and Integrative Genomics Viewer, IGV.

### Data intersection

Intersection between pairs of the genomic regions was performed using bedtools intersect (v2.19.1) ^[53]^.

## DATA AVAILABILITY

Raw and processed Hi-C, ATAC-seq, and RNA-seq data from this study are available from the NCBI Gene Expression Omnibus as series GSE126117.

## SUPPLEMENTARY FILES

**Figure S1.**
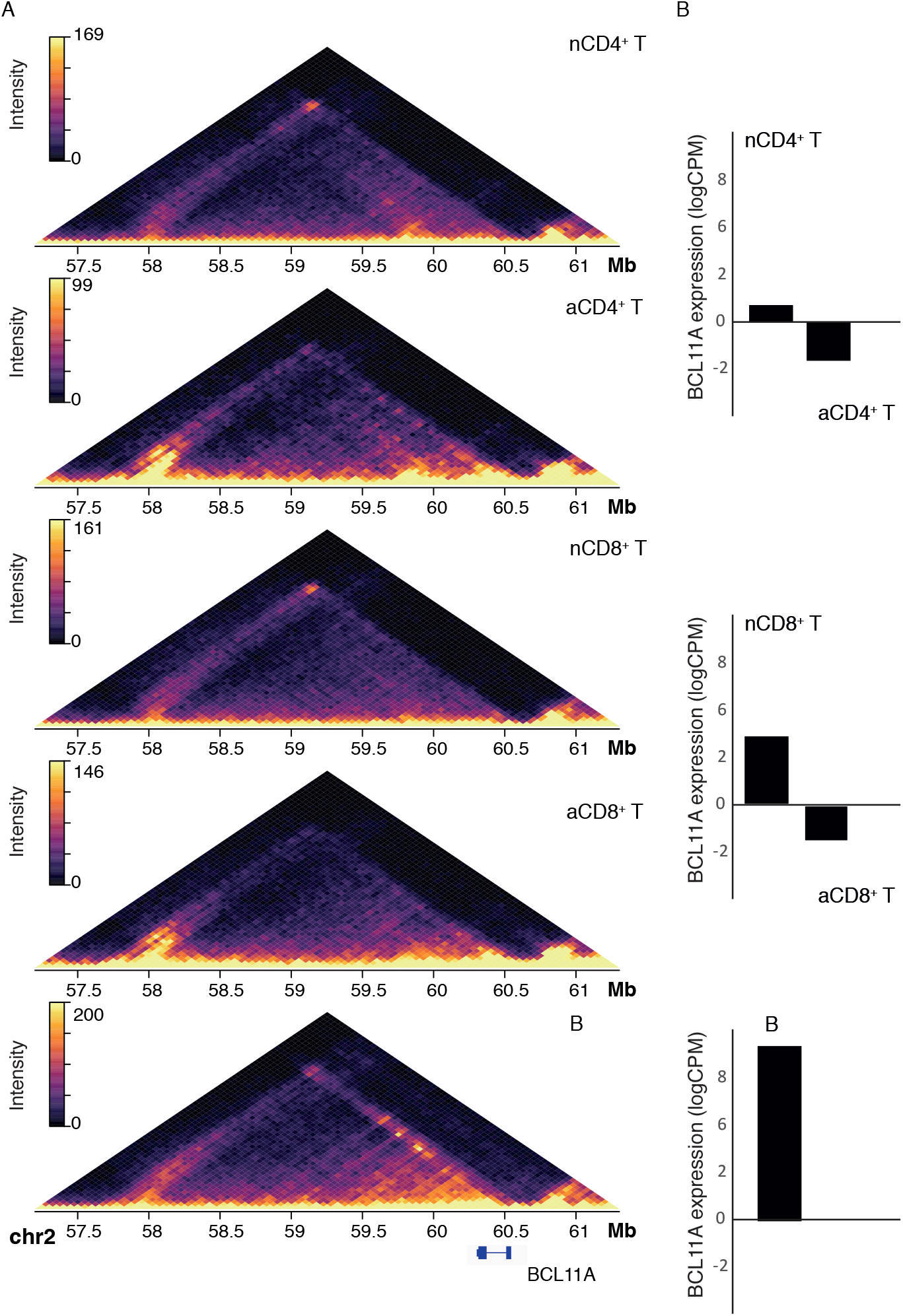
Chromatin structure and gene expression are immune cell type-specific. (A) *In-*situ Hi-C contact matrices for a 2.5 Mb region on chromosome 2 that includes the BCL11A locus at 50 kbp resolution. The top four Hi-C matrices display data from resting and activated T cells, while the bottom Hi-C matrix shows data from B cells. Color scale indicates number of reads per bin pair. (B) Bar plots showing the expression of BCL11A across resting and activated CD4^+^ and CD8^+^ T cells, and B cells.

**Figure S2.**
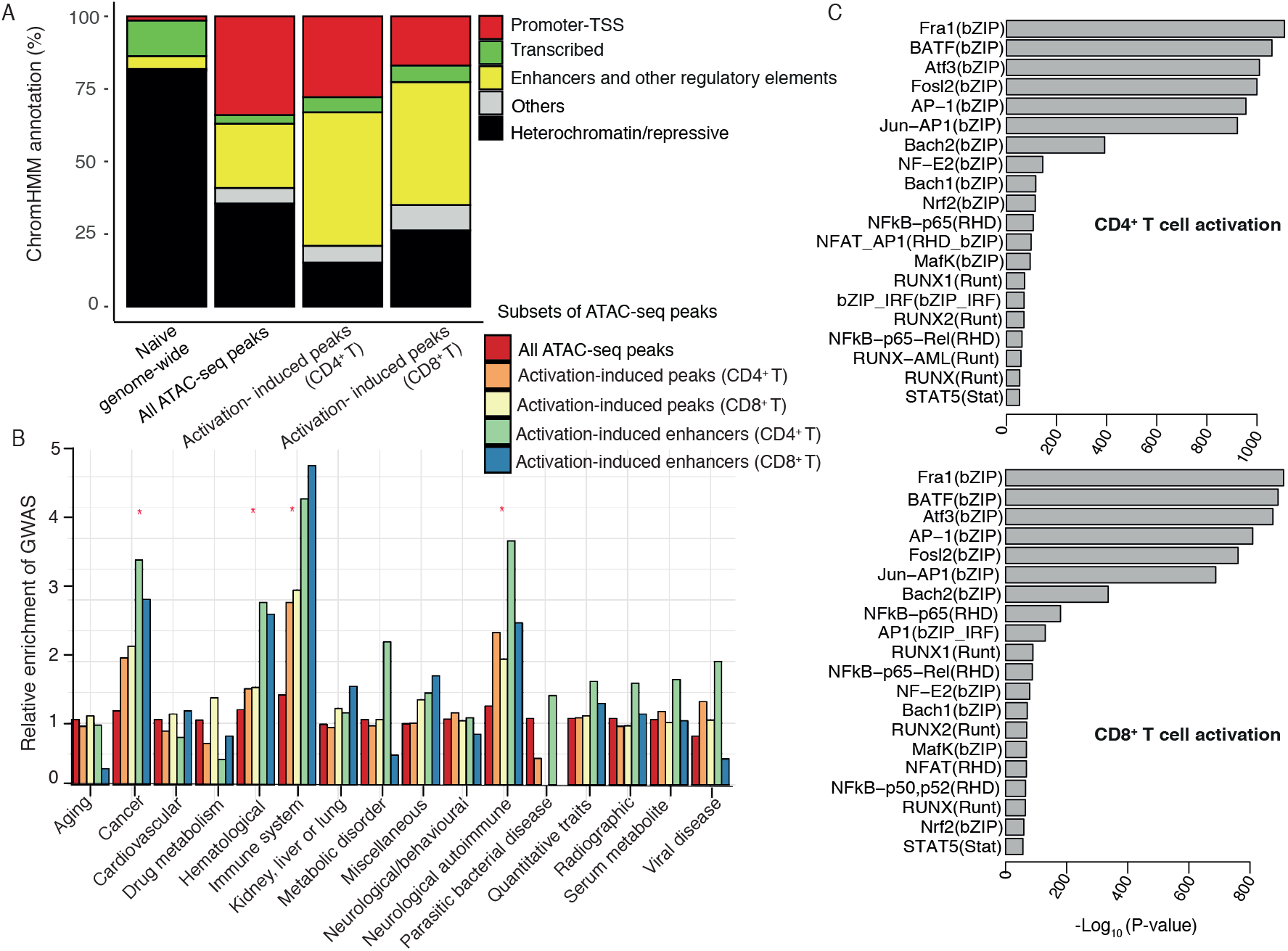
T cell activation-induced accessibility changes are enriched for enhancers and other regulatory elements and GWAS SNPs. (A) Bar plots showing the distribution of the predicted chromatin states (ChromHMM) in subsets of peaks. (B) Bar plots showing enrichment of GWAS SNPs in subsets of ATAC-seq peaks. Shown are the relative enrichment of GWAS (observed/ expected number of overlapping SNPs) (Y axis) for each SNP category (X axis). Red asterisks denote SNP categories for which subsets of ATAC-seq peaks showed significant enrichment by GREGOR.(C) Bar plots showing the top-ranked TFBS motifs that are enriched in activation-induced accessibility changes in CD4^+^ and CD8^+^ T cells, where the X-axis represent the significance of the enrichment (−log10 P-value, cumulative binomial test), and Y-axis each of top-ranked motifs.

**Figure S3.**
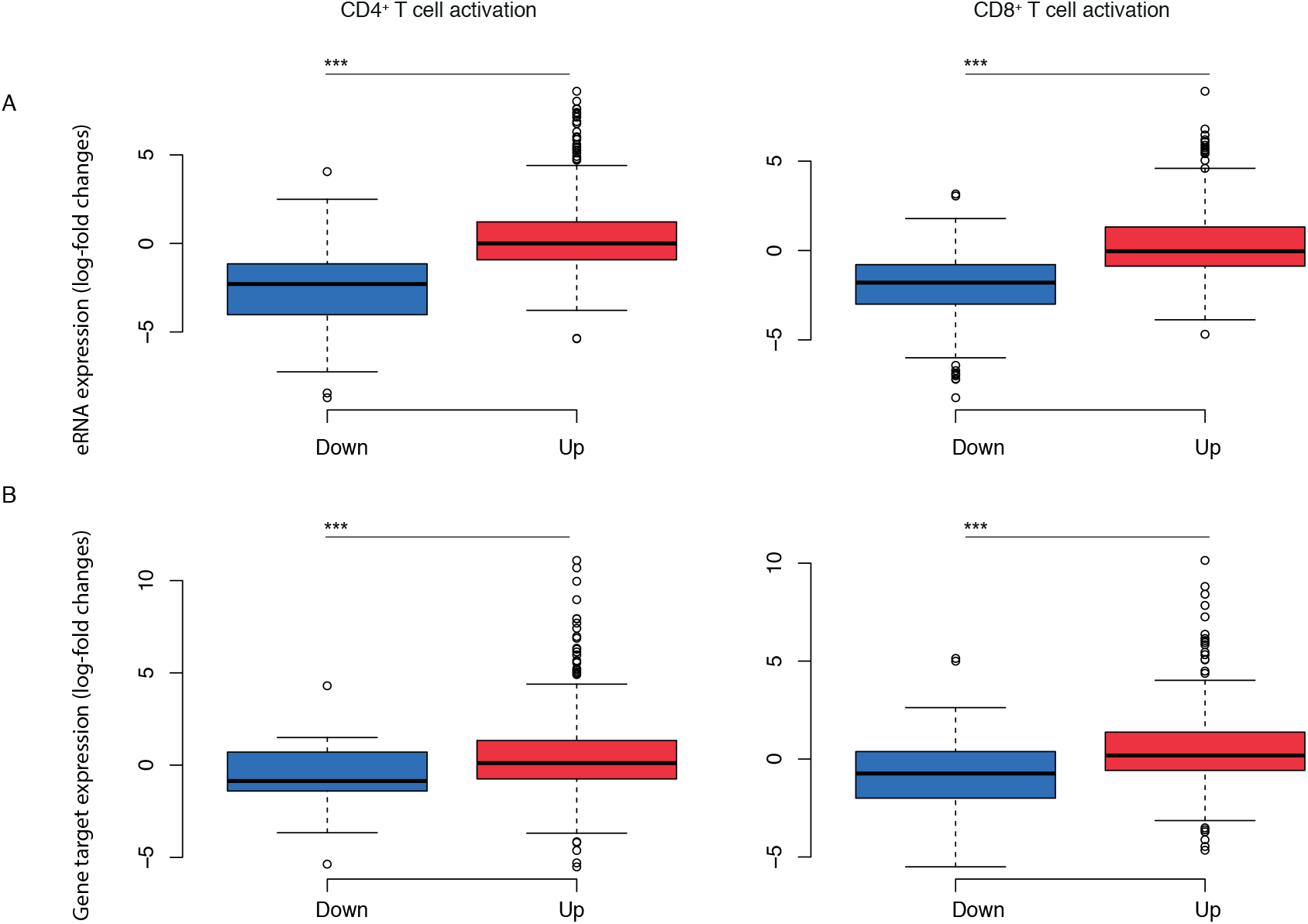
Variation of chromatin accessibility at the activation-induced peaks is associated with changes in expression of eRNAs and targets genes. Distribution of the expression of (A) eRNAs and (B) predicted enhancer target genes across the activation-induced peaks that gained and lost accessibility is represented as box plots (central bar represents the median with boxes indicating interquartile range, values of the upper and lower quartiles are represented in with whiskers and outliers with circles). Statistical comparisons were made with the unpaired Wilcoxon test: ***P-value < 0.0001.

**Figure S4.**
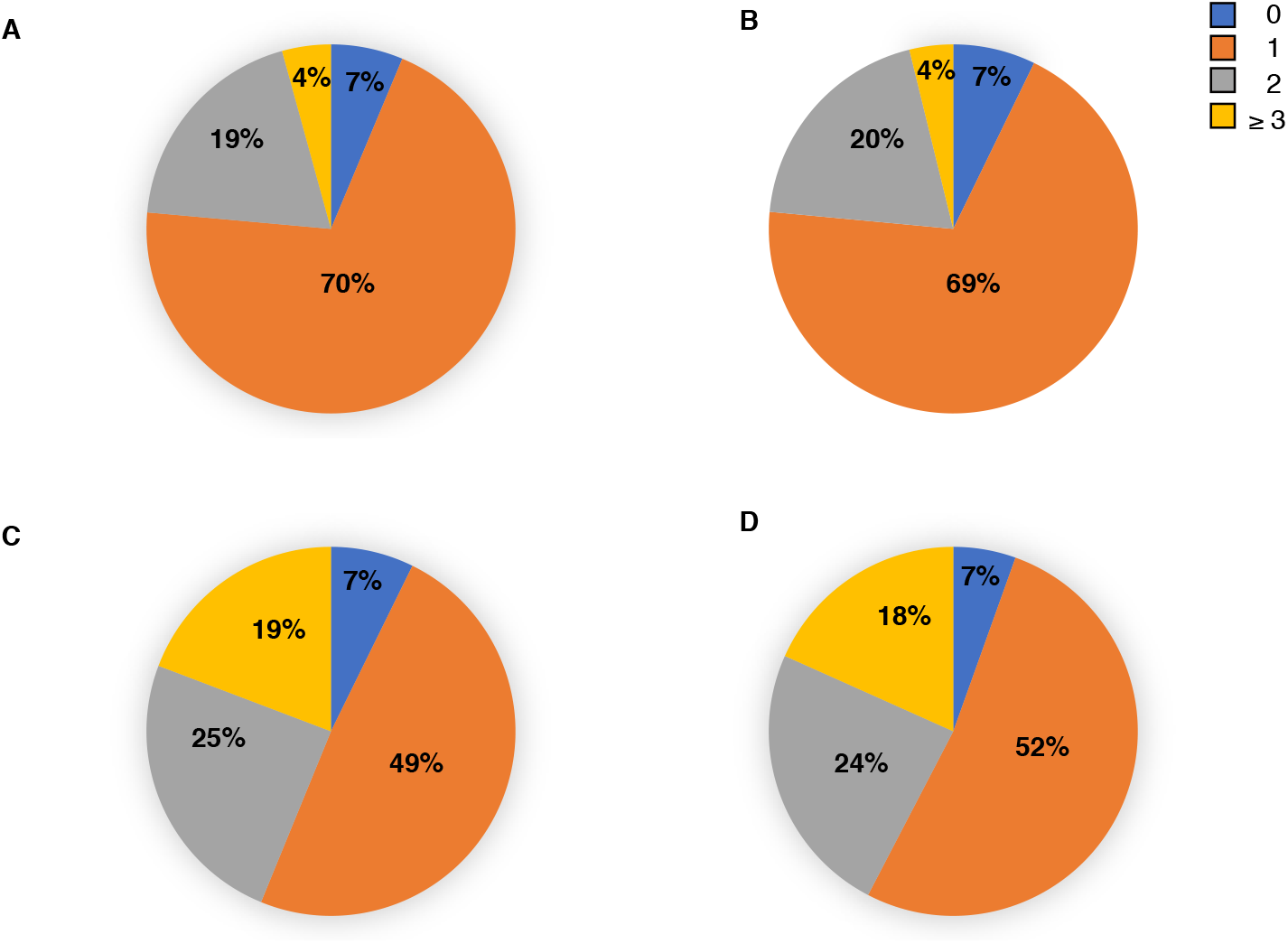
T cell activation results in partitioning of genome topology. Pie chart representation of the distribution of (A) TADs in resting CD4^+^ T cells overlapping 0,1, 2 or ≥ 3 TADs in resting CD8^+^ T cells, (B) TADs in resting CD8^+^ T cells overlapping 0,1, 2 or ≥ 3 TADs in resting CD4^+^ T cells (C) TADs in resting CD4^+^ T cells overlapping 0,1, 2 or ≥ 3 TADs in activated CD4^+^ T cells, and (D) TADs in resting CD8^+^ T cells overlapping 0,1, 2 or ≥ 3 TADs in activated CD8^+^ T cells.

**Figure S5.**
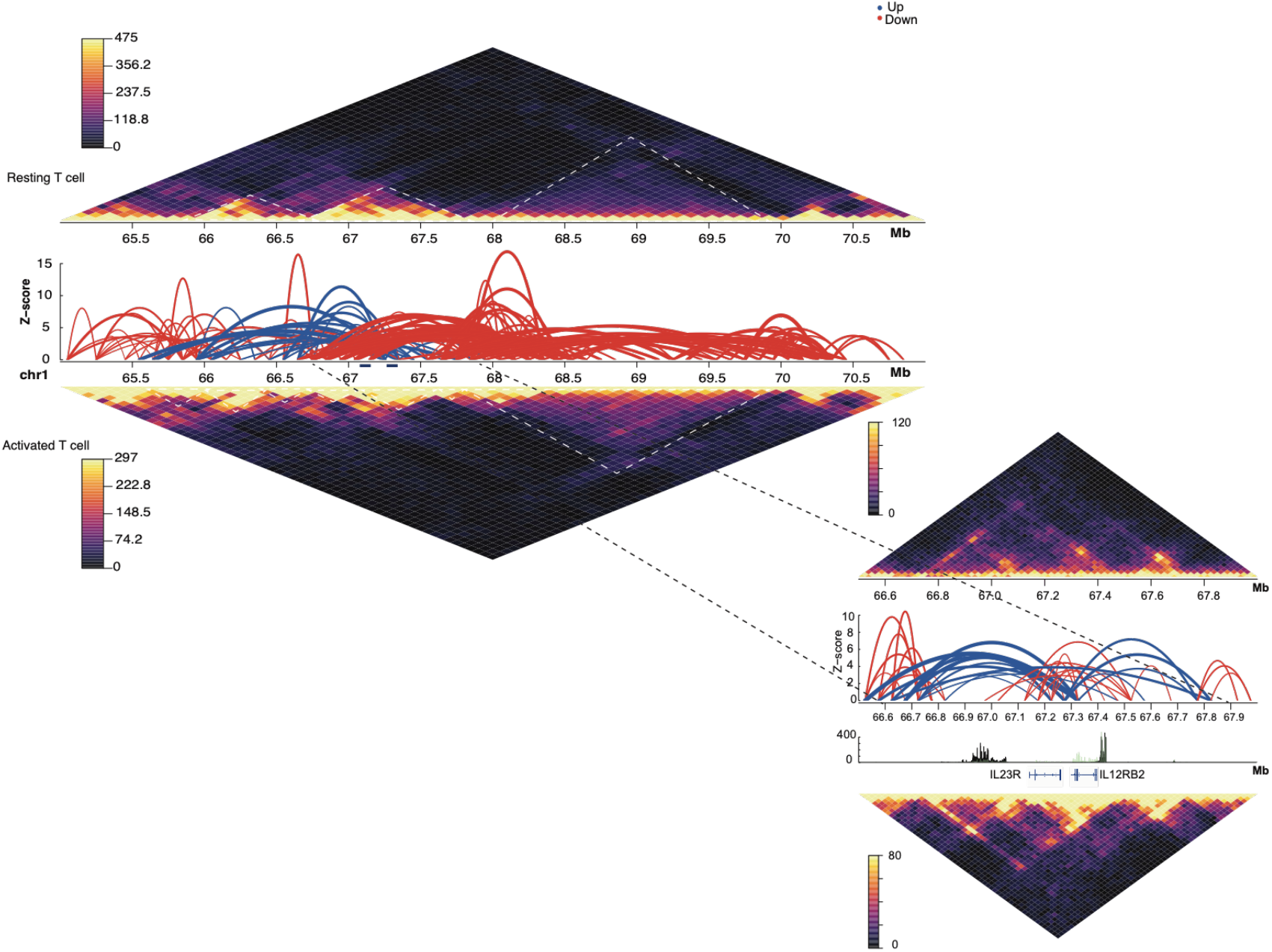
T cell activation results in partitioning of genome topology. In-situ Hi-C contact matrices at 100kbp in resting and activated T cells over genes that have been associated with T cell activation such as IL23R and IL12RB2. Colour scale indicates number of reads per bin pair. Locations of the TADs as called by TADbit at these regions are indicated below each contact matrix. Significant differential interactions at 100 kbp (FDR<0.05) are represented by arches where the z-score is −log_10_(p-value). Red and blue lines represent gain and loss of interactions, respectively, in response to activation. The zoomed inset regions show in-situ Hi-C contact matrices, significant differential interactions at 25 kbp resolution (FDR<0.05) and RNA sequencing coverage plots for the IL23R and IL12RB2 loci. Turquoise and black coverage plots represent activated and resting T cells respectively.

**Table S1: Differential accessibility analysis comparing naïve to activated CD4^+^ and CD8^+^ T cells**.

**Table S2: List of disease- and trait-associated variants**

**Tables S3A and S3B: (A) Differential interactions comparing naïve to activated CD4^+^ (A) and CD8^+^ (B) T cells**.

